# Existence and construction of large stable food webs

**DOI:** 10.1101/097907

**Authors:** Jan O. Haerter, Namiko Mitarai, Kim Sneppen

## Abstract

Ecological diversity is ubiquitous despite the restrictions imposed by competitive exclusion and apparent competition. To explain the observed richness of species in a given habitat, food web theory has explored nonlinear functional responses, self-interaction or spatial structure and dispersal — model ingredients that have proven to promote stability and diversity. We here instead return to classical Lotka-Volterra equations, where species-species interaction is characterized by a simple product and spatial restrictions are ignored. We quantify how this idealization imposes constraints on coexistence and diversity for many species. To this end, we introduce the concept of *free* and *controlled* species and use this to demonstrate how stable food webs can be constructed by sequential addition of species. When we augment the resulting network by additional weak interactions we are able to show that it is possible to construct large food webs of arbitrary connectivity. Our model thus serves as a formal starting point for the study of sustainable interaction patterns between species.

## Introduction

When interactions between species are random, stability of large complex ecosystems has been suggested to be compromised [1]. On the other hand, when all interactions are absent, populations grow exponentially and diversity is suppressed since the winner will “take all”, i.e. the most-competitive species will take over all available resources. These considerations imply that neither the presence of random interaction nor its complete absence can explain the observed diversity and apparent stability. It may thus be non-random competition between species that is needed to explain the diversity of the living word.

One step towards better understanding of how interactions can stabilize ecosystems comes from the web formed by phage and bacteria species in the ocean, where the dominance of the fastest growing bacteria species is reduced by phage predation [2]. Using oceanic data [3, 4] we recently demonstrated that the species richness of bacteria is strongly correlated with similarly high species richness of their phages, a feature we described as a “staircase of coexistence” [5]. This phenomenon of mutual support of diversity reflects a generalized version of competitive exclusion [6]. Notably, the resulting ecosystem has non-random interaction strengths, a manifestation of the principle of trade-offs [7, 8, 9].

As mentioned above, R. May [1] called into question whether well-connected food webs could be dynamically stable—using the assumption of random interactions. For observed food webs, however, it has by now repeatedly been stated that interactions between species populations are far from random [10, 11, 12, 13, 14, 15]. A body of experimental [16, 17, 18] and theoretical work [19, 20] suggests, that the interaction-strength distribution may be strongly skewed, with only few strong, but many weak interactions. It was further emphasized that the patterning of links might play an important role in stability [10], contradicting the null model of purely random interactions. The stabilizing role of non-randomness is further corroborated by analysis of loops [11]. Neutel and collaborators found that real food webs are structured such that long loops contain more weak links than expected at random, thereby promoting stabilization. Realizing that even May’s random matrices implicitly make assumptions on the frequency of interaction types, recent work on dynamic stability stresses that more realistic relations between parameter values should be incorporated [14, 15, 21]. Their conclusions are surprising, in that they associate realistic link structures with lower stability than their unstructured counterpart, a finding yet to be reconciled with earlier studies [22, 23].

When exploring the stability of a specific fixed point for a given food web, the existence of this fixed point is implicitly assumed. However, the available theoretical criteria for dynamical stability so far remain difficult to reconcile with feasibility [21], defined as the demand that all species have positive populations. In the present work, we depart from the quest for random interactions and instead explore possible patterns of interactions that allow for large food webs under the strict constraints on both feasibility and stability.

Here we return to the classical Lotka-Volterra equations, where interactions between consumers and resources are modeled by the linear type-I response, i.e. the rate of consumption is proportional to the product of concentrations of the consumer and its resource. We first review the food web assembly rules, which describe how such equations yield a linear set of equations for the steady-state solutions of species concentrations and how such linearity constrains possible link structures in multi-level food webs (Methods). This leads us to introduce the concept of *free* and *controlled* species. For the subset of network structures compatible with the assembly rules we then describe how these could be built-up systematically, starting from a single resource and demanding feasibility and global stability at every addition of species. We further demonstrate how sizable communities of dozens of species can be obtained by this methodology and discuss the role of consumption efficiency, i.e. the rate of using resource biomass for growth. We finally compare species richness in our constructed communities to that of field data and close with a discussion of the implications of our work.

## Methods

### Generalized Lotka-Volterra equations

In the web formed by the interactions between consumers and resources in a habitat, energy flows from a physical or chemical nutrient source up the food chain. The resulting structure is a food web which ties together the resources, their consumers and subsequent consumers. Consider first a food web of *L* trophic levels (Fig. 1a) where each species can be assigned a sharp trophic level *l*, i.e. any species exclusively consumes other species at a specific level. Such webs have recently been termed *maximally trophically coherent* [21]. Using Lotka-Volterra-type formalism, with the implicit assumptions of a well-mixed ecosystem governed by mass action kinetics, the energy flux passing through a species 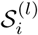 at level *l* is [24, 25]

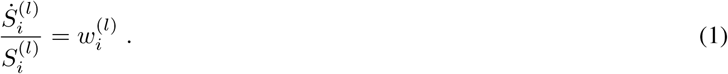

Here and throughout the text use the symbol 𝒮 as a general label for a given species, while *S* and *Q* refer to its time-dependent and steady-state concentrations, respectively. When formulating the RHS of Eq. 1 in units of the carrying capacity for primary producers (*l*=1), the flux

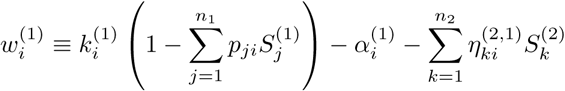

incorporates logistic growth at the basal level whereas

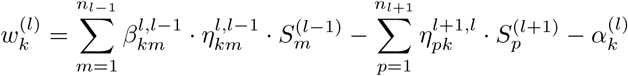

accounts for biomass conversion at higher levels *l* > 1. Here, 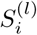 is the biomass concentration of species 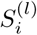, superscripts (subscripts) label the trophic level (species within a trophic level), 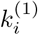 are maximal growth rates of basal species and *p_ji_* are the basal nutrient depletion strengths, which we subsequently will set equal to unity.

**Figure 1:**
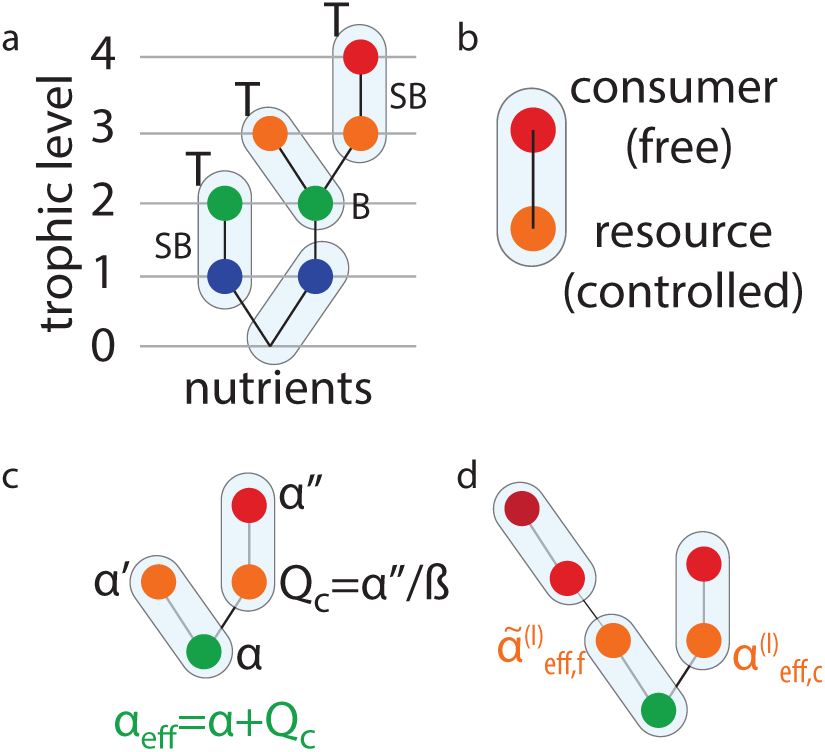
Definitions. **a**, Simple food web consisting of four trophic levels. We define the nutrient source to be trophic level zero. The trophic level of any species is the average trophic level of its resources plus one. Colored circles mark species, light blue ovals mark pairings of species, symbol (T) marks top species, i.e. species without consumers. Side branches are marked by the symbol “SB”. A branching species (or branching point) is marked by “B”. *Q_c_* = *α*″/*β* **b**, An example of a pairing, consisting of a “free” consumer and a “controlled” resource species. **c**, Definition of *α_eff_* for a controlled species: *α_eff_* equals the sum of the decay coefficient of the species itself and the steady-state density of its controlled consumer(s). **d**, Illustration of the condition (Eq. 10) for feasibility at branching points, showing the comparison between 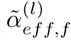 and 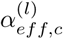 for two branches formed by a free, respectively controlled, species.

In the above equations we use the rate 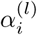 as the species-specific decay coefficients of species *i* at trophic level *l*, whereas 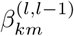 is the link-specific consumption efficiency of a species *k* at level *l* when consuming the species *m* at level *l* − 1, and 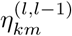 are the corresponding link-specific interaction strengths. For species at the top level *L*, the loss term in 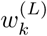 will only contain the intrinsic decay rate 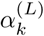 representing death by aging, accidents, or disease. For later use, independent of their trophic level, we call species without consumers *top species.*

### Assembly rules

In steady state, 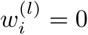 and Eq. 1 is a set of linear equations. A necessary condition for a unique steady-state solution is nonzero determinant of the interaction matrix, which leads to the recent food web assembly rules [26]. Uniqueness in turn is required for structural stability of the solutions. The assembly rules break the food web down into *pairings* of species. A pairing consists of a consumer and one of its resources, hence the two species of each pairing are connected by an existing feeding link (Fig. 1a,b). The existence of a steady state in a maximally trophically coherent food web strictly requires that the food web can be covered by such pairs and none of these pairs overlap [26]. This is known as *perfect matching* in graph theory [27]. Physical or chemical nutrients can, but need not, be part of such pairings.

The pairing is convenient as it guarantees that no group of two or more species share exactly the same niche, i.e. a particular set of interactions. The resulting rules can be seen as a generalization of the competitive exclusion principle, which states that when two consumers compete for the exact same resource within an environment, one consumer will eventually out-compete and displace the other [6, 28, 29]. Considering the different nutrient sources, e.g. sunlight or chemical substances, as residing at trophic level zero (Fig. 1a), the assembly rules [26] can be stated compactly:

*The number of species at any trophic level l must not be less than the number of free species at the level l* + 1.

## Results

In the following we introduce the conceptual classification of species into free and controlled species. Using these concepts, we then provide conditions for sustainable additions of species, i.e. conditions that allow both the resident and “invading” species to coexist without any species becoming extinct. These conditions relate the intrinsic fitness of an “invader” to that of a specific subset of species providing for the invader's nutrient uptake. We find that the division into “free” and “controlled” species is natural in defining such conditions. Originating from this methodology, we then discuss how sizable food webs can be constructed. Finally, we compare the species richness in our synthetic food webs to that observed in the field.

### Free and Controlled Species

For the subsequent discussion, we define a class division of species into either the “free” or the “controlled” categories. Free species are either top species, or species which are exclusively predated upon by controlled species. In contrast, a controlled species is one that is regulated by (exactly one) free consumer. Note that the case of two free consumers is ruled out by competitive exclusion. Besides this free consumer, a controlled species may be preyed upon by any number of controlled consumers. The motivation for this division into “free” and “controlled” is a basic consequence of type-1 interactions, namely that a free predator sets the population size if its prey, irrespective of the prey’s reproduction rate. For simple linear food chains or for hierarchical food webs each species is uniquely determined to be either free or controlled, reflecting the uniqueness of non-overlapping pairing [26] in such networks.

### Species addition

Consider now the process of adding new species to an existing food web. Following the addition of a few individuals of the new species, Eq. 1 describe the subsequent dynamics of species populations towards a new steady state “equilibrium” value. We will now explore conditions for stability and feasibility of the steady state for the thereby obtained new food web. Throughout, we will assume a separation of the fast timescale of dynamical adjustment of all species concentrations to an equilibrium value and a slow timescale of the introduction of any new species. We restrict additions to occur on a one-by-one basis. For simplicity, we will refer to newly added species as invaders, but note that they might more properly be referred to as “additions” or “introductions“, since their properties will be chosen as to allow for coexistence of the resulting food web.

#### Conditions for stability

A sufficient criterion for dynamical stability is the existence of a Lyapunov function [24, 30]. Assume first that positive, i.e. feasible, steady-state biomass concentrations 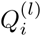 exist for all species. 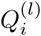 is then obtained from setting 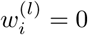 in Eq. 1. In analogy to Goh [30], we define the function

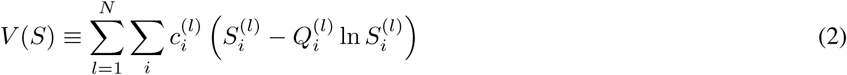

with constants 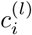. The time derivative of this function is

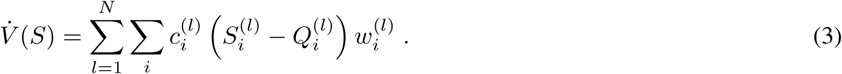

Inserting 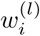 from Eq. 1 into Eq. 3 one obtains sums of fluctuations of species concentrations. When demanding that *V̇*(*S*) be strictly negative, conditions result which relate the consumption efficiencies to the constants 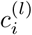. For *l* > 1 and *l*=1 the respective conditions are [26]

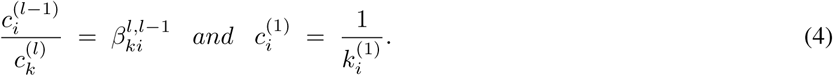

Such *c*’s can always be found if each species is only connected to the basal level by one path, as any *c* parameter then can be chosen as a simple product of the *β*’s present along the path to the basal level. In the more general case where some consumers feed on two prey species, the constraint will typically be violated, and if the product of *β*’s along different paths differs by a large factor the food-web might become unstable. In the case of omnivorous consumption such alternative weighted paths may even cause chaos [31].

Notably, Eq. 4 constrains 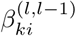 and 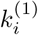, but is independent of the interaction strengths 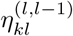 and decay rates 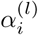. Stability is hence unaffected by these parameters and they can therefore be adjusted independently to achieve feasibility. As a noteworthy side remark, when defining the total and relative steady-state biomass, *Q_tot_* ≡ ∑_*i*_ *Q_i_*, respectively *q_i_* ≡ *Q_i_*/*Q_tot_*, one can express the steady state value of the Lyapunov function as *V*(*Q*) = *Q_tot_* [1 + *ln D* − *ln Q_tot_*]. Remarkably, this function only depend on *Q_tot_* and the diversity *D* ≡ exp (− ∑_*i*_ *q_i_ln q_i_*).

When the assembly rules [26] are fulfilled, a non-overlapping pairing of species can always be found. When only the links involved in pairs are considered, and the resource species of each pair is connected to another species/nutrient at the trophic level below, the resulting graph is a tree. In such a tree, or equivalently a hierarchical food web, each species can consume only one other species, or the nutrient source. These tree-like webs are particularly convenient, and minimal in the sense that each species contributes only one link. They constitute an extreme case of trophically coherent networks, as all species trivially have a unique trophic level.

The tree-structure generalizes the recently noted two-branch-structure [32], where rules for the three cases of even-even, odd-even, and odd-odd branches were described. As indicated above, the relation between 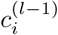 and 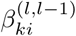 can always be fulfilled by appropriate choices of 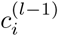 in a tree-like food web, without even making any assumptions on 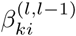. Thus stability can be obtained, irrespective of the choice of any of the parameters for the link strengths or decay coefficients.

#### Conditions for feasibility

Although dynamical stability for tree-like food webs can always be achieved, it gives no answer to the existence of feasible steady state densities *Q_i_* > 0. To prove that feasibility can be obtained, we consider the steady state of tree-like food webs and for simplicity specialize to 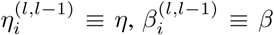, and 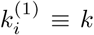, for all links, i.e. we assume all species to have similar interaction and growth coefficients. The species specific decay coefficients 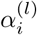 are free parameters. We will show that even under these constraints, positive steady state solutions can always be obtained.

In steady state *w^(l)^* = 0 in Eq. 1 allows us to determine all populations in a tree-like food web uniquely. Dividing the equations 1 through by *η*, this interaction strength only appears in terms of the form *k*/*η* and 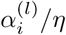. For simplicity we hence absorb *η* in these two symbols, i.e. *k* → *k*/*η* and 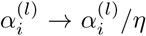. The equations separate species into two types, also discussed in the method section:

- For the first type the species biomass concentrations are entirely determined by sums of decay rates higher up in the food chain. These are the *controlled* species characterized above (Fig. 1b).
- The second type are *free* species, which have concentrations that are also affected by species or nutrients lower in the food chain (Fig. 1b). In simple food chains, such species are sometimes called *food limited* [32].

As mentioned above, a controlled species has exactly one free consumer. Further, both free and controlled species can have any number of controlled consumers.

Using Eq. 1 we find the steady state biomass concentrations of a free species at a trophic level *l* ≥ 2

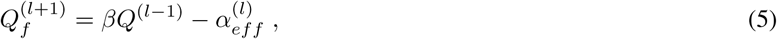

where unnecessary subscripts were dropped for simplicity. Equation 5 expresses the biomass concentration of a free species in terms of resources available to its prey (thus at level *l* — 1) minus resources that are consumed because its prey is exposed to various other sources of death. This include both spontaneous death, as well as possible predation by other consumers, described through the effective decay rate

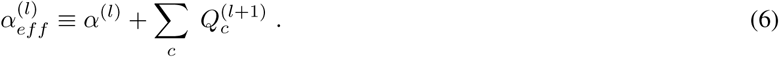

The quantity 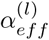 thus include intrinsic decay plus decay caused by all its consumers that are controlled from above, see Fig. 1c. Thus controlled consumers contribute to the effective decay rate of their resources, whereas species without controlled consumers only are exposed to their own limitations. Notice that we persistently use the subscripts *f* and *c* to denote free and controlled consumers of the corresponding species, whereas absence of subscript implies that this species may be of either of the two types.

Let us for example consider the simple case where 𝒮^(*l*)^ is the biomass concentration of a top consumer that preys on a resource 𝒮^(*1*−1)^. In that case 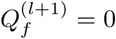 and Eq. 5 simplifies to 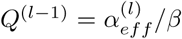. Thus, the biomass *Q*^(*l*−1)^ is proportional to the decay rate of its single free consumer including the loss to whatever that eats this consumer (Eq. 6). Because each species can act as a resource for at most one free consumer (competitive exclusion), these other predators must all be controlled from above.

Equations 5 and 6 allow us to determine all steady-state populations in a tree-like food web. When *Q*^(*l*−1)^ is free, an iterative loop unfolds by Eq. 5, until a species with known density or controlled density is reached somewhere at a lower point in the food-web. In some cases, this may demand continuation of the iterative loop (Eq. 5) until the basal level (*l* = 1) is reached. Closure of the system using the nutrient source similarly involves the interplay between free and controlled species at the basal level.

As for the species, the nutrient source may be either free or controlled. For free nutrients (Fig. 2a), all basal species are controlled and their densities 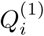 are the (known) effective decay coefficients 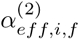 of their respective free consumers 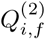. The steady-state equation for basal species (Eq. 1) for each 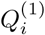 gives the population densities of the free consumers of 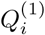:

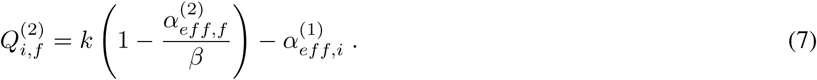

where 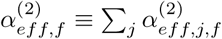 is the sum of all effective decay rates of the controlling (free) consumers at level two.

**Figure 2:**
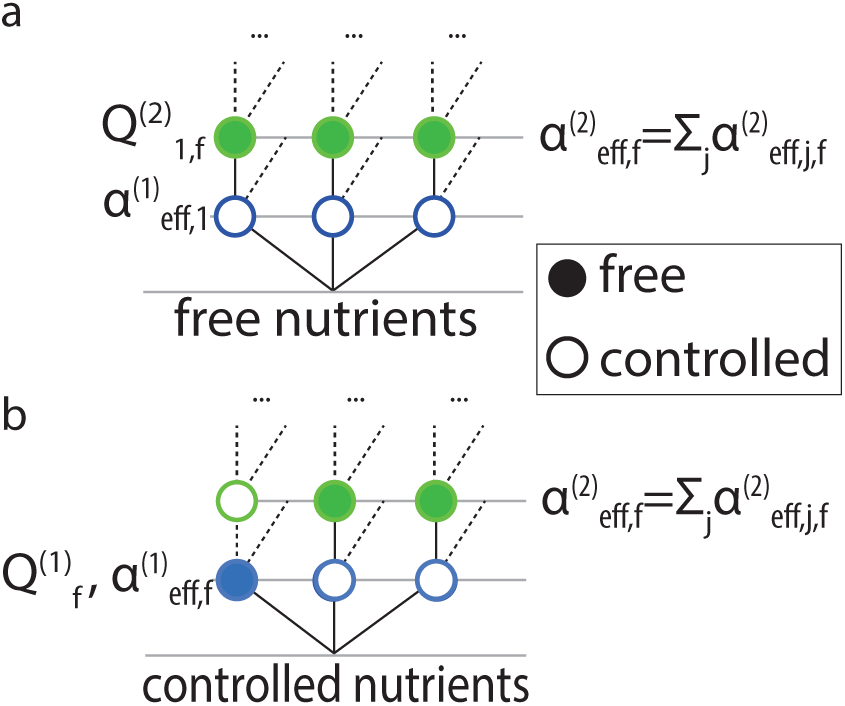
Closure at the basal level. **a**, Case of free nutrients: All basal species are controlled. **b**, Case of controlled nutrients: One basal species (density 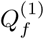) is free. Dashed lines in both panels symbolize links to possible controlled consumers.

For controlled nutrients (Fig. 2b), one basal species is a free consumer of nutrients. Its population density equals

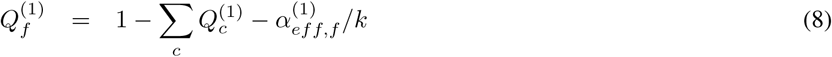

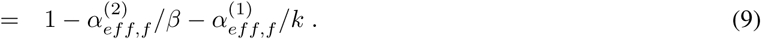

Using either Eq. 7 or Eq. 9 in Eq. 5, the remaining free species densities can be determined.

When *Q*^(*l*−1)^ is controlled, Eq. 5 determines a species’ population from its controlling consumers and their consumers further up in the tree. At the same time, *Q*^(*l*−1)^ is determined by the effective decay rate of the corresponding free species 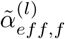 as 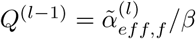 (see Fig. 1d). Since this determines 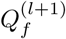 by Eq. 5, the constraint 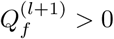 amounts to

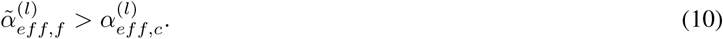

The constraint in Eq. 10 specifies an inequality at each branching point of the tree-like food web. It amounts to saying that the effective decay rate along the free branch has to be sufficiently high, in order to maintain the controlled branch. If the free branch is more competitive than this limit dictates, then the controlled branch will at least partially collapse.

### Construction of food webs

Given sustainability in terms of stability and feasibility we now systematically build up food webs. By this we mean that species are attached one-by-one to an existing sustainable food web in a way that the resulting food web again is sustainable. When composing food webs by sequential addition of species, we first note that for tree-like food webs, any added species will be a free consumer. This means that it will be controlling its resource, and if also another species controls this resource, competitive exclusion dictates that the one with the smallest effective decay rate will either eliminate the other or an odd number of consumers, until the condition in Eq. 10 is again met. A successful invader will thus be free, and all other consumers of its resource must be controlled. Further, the invader has to fulfill Eq. 10, implying that it must not be too efficient. This amounts to a lower bound on an invading species' decay coefficient in order to maintain feasibility for existing species. An upper bound is given by the requirement that also the new species should be feasible.

When inserting a new species at a small concentration, successful entry amounts to its decay coefficient being sufficiently small. E.g. for an initial basal species this requirement is simply that *α*^(1)^/*k* < 1. Adding a second species at trophic level two, the condition that *Q*^(2)^ should be positive gives *α*^(2)^/*β* − *α*^(1)^/*k* < 1, a criterion that is easier to meet for large consumption efficiency *β*. To exemplify assembly of a sizable food web, we coarse-grain decay coefficients into steps of 1 ≫ *α_min_* = 0.02. Requiring this value also as a lower bound, Eq. 10 implies a sometimes substantial *α*, especially for those species that are added to larger food-webs.

Our assembly starts with one basal species, and proceeds by iteratively making random additions of species as consumers to any existing species. An addition is only accepted whenever the assembly rules allow coexistence of both the invader and the previous species. Thus our assembly is not an attempt to mimic evolution, but rather an idealized procedure to construct stable and feasible food webs.

Note that in tree-like food webs, the trophic level of any particular species is determined by its single resource. We now set *k* = 1 in the remaining discussion as well as the simulations in Figs 3—4. The first species addition (Fig. 3a—c) yields a biomass density 1 − *α_min_* and the added species is part of the first trophic level. As illustrated in Fig. 3a, subsequent additions can lead to rather large food webs. Panel b illustrates how the population sizes are distributed as the food web grows. One observes that for small systems populations are either small or large, whereas larger food webs have a single-peaked distribution of biomass (also quantified in Fig. 3e).

**Figure 3:**
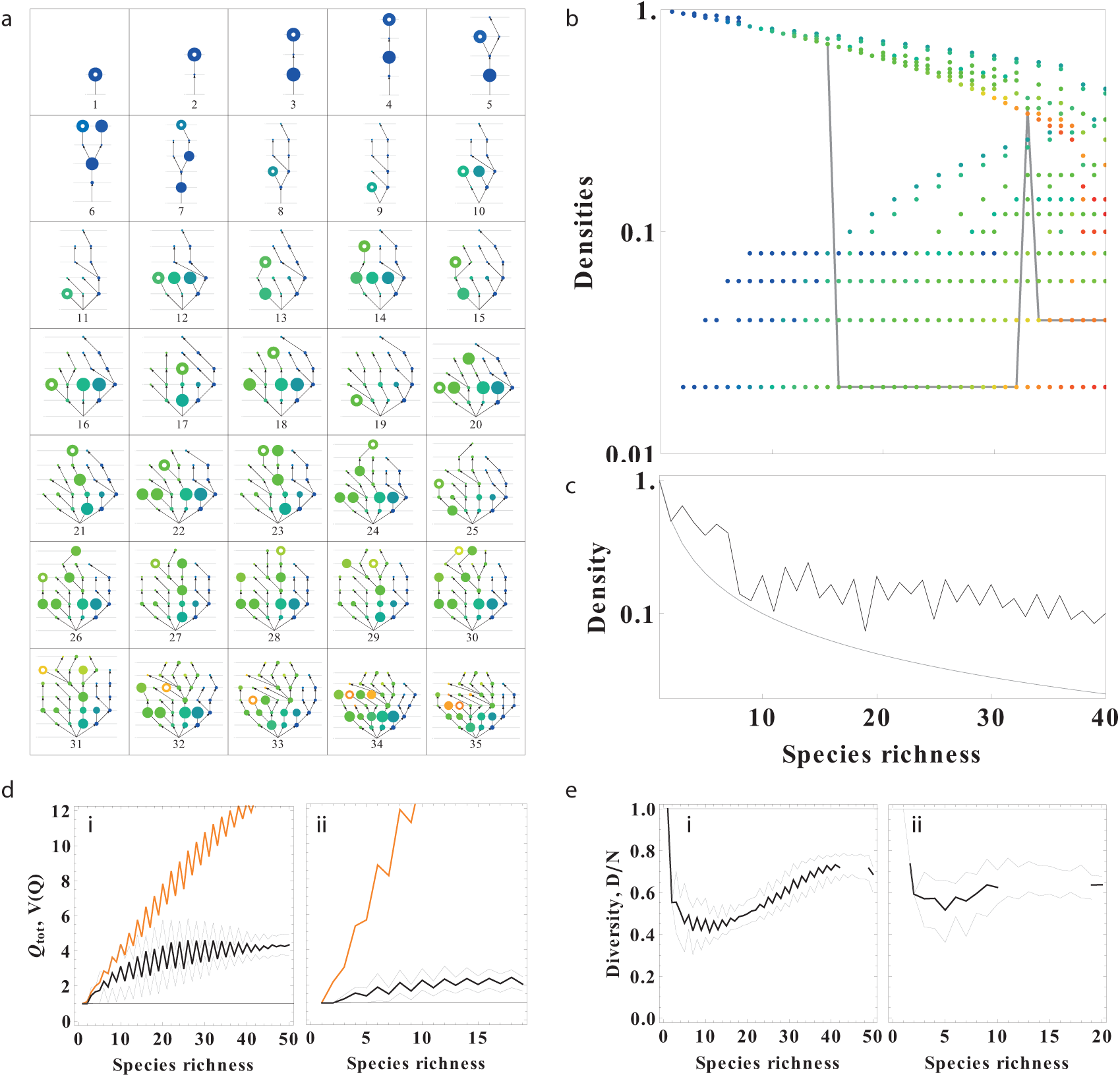
Assembly of a sustainable model food web. **a**, Sequential additions, starting from a single basal species. Colors from blue to orange indicate early and late additions, respectively. Area of nodes is proportional to steady-state species densities *Q_i_*. Lines indicate consumer-resource links. Latest addition is marked by a white point in each panel. Number of addition marked below the webs. Note that after each addition, a new non-overlapping pairing (Fig. 1) can be found. **b**, Steady-state concentrations *Q_i_* vs. species richness for the web in (a), in correspondingly chosen colors. Gray line: trajectory of a particular species, addition 15, compare (a). **c**, Average concentration 〈*Q*〉. Smooth line indicates the inverse of species richness. Note the logarithmic vertical axes in panels (b) and (c). **d**, Total biomass (black) and Lyapunov function (orange) averaged over multiple realizations of food webs as a function of species richness. Thin lines indicate the 10th and 90th biomass percentiles, respectively. **i**, *β* = 1, **ii**, *β* = 1/2. **e**, Diversity *D* = exp (− ∑_*i*_ *q_i_ln q_i_*) per species for *β* = 1 (**i**), respectively *β* = 1/2 (**ii**).

**Figure 4:**
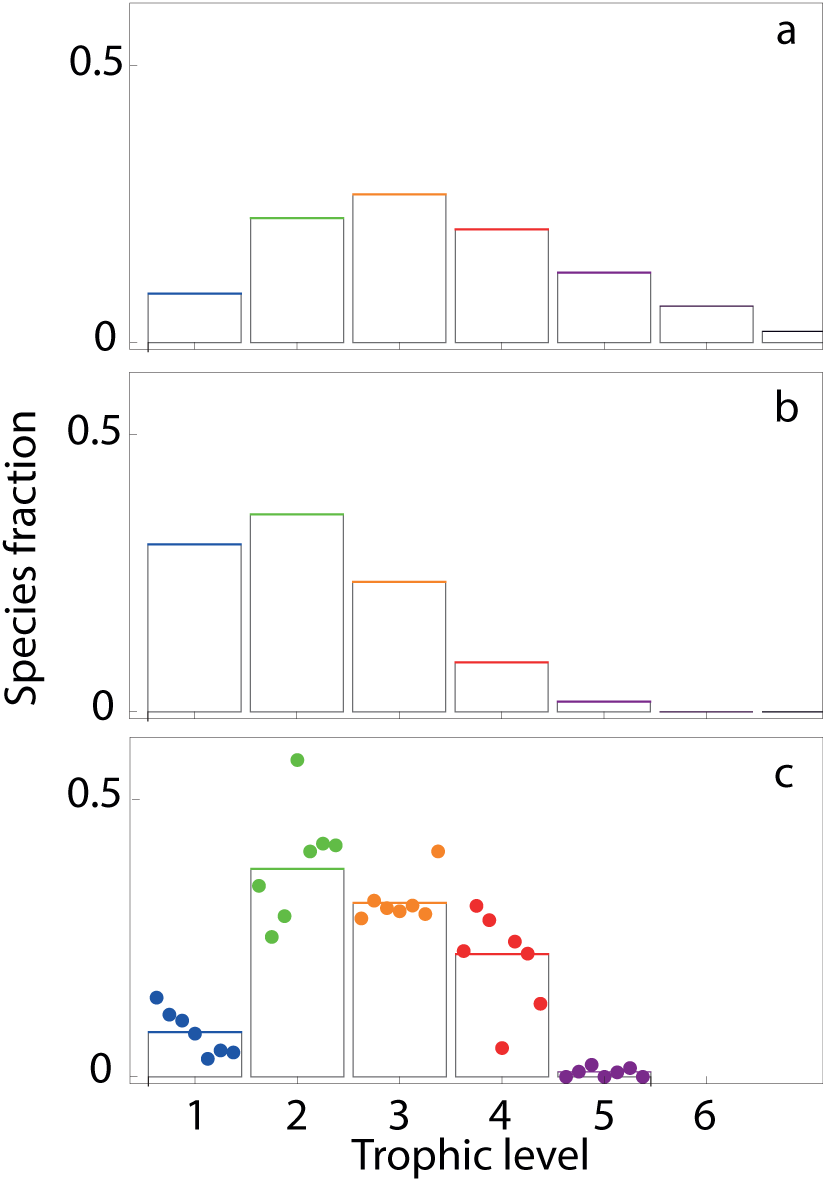
Distribution of species richness according to tropic levels. **a**, assembly prediction with *β* = 1 and *α_min_* = 0.01. **b**, assembly prediction with *β* = 0.2 and *α_min_* = 0.01. **c**, empirical data from 7 different food webs, compiled by Dunne et al. [33, 34]. The original data are: Carpinteria Salt Marsh, Estero de Punta Banda, Bahia Falsa[35, 36]; Flensburg Fjord [37], Sylt Tidal Basin [38]; Otago Harbor [39]; and Ythan Estuary [40]. The trophic level of a given species was determined as the average path length to the nutrient level when following predation links downwards towards the nutrient level.

With every species addition, transitions of biomass occur, with invaders typically starting with relatively high biomass. The solid line in panel Fig. 3b illustrates that the addition of a species may cause drastic changes in populations of resident species. Note that this change is associated with a shift from acting as a controlled to acting as a free species. This can e.g. be seen in Fig. 3a, addition 5, where an added consumer reduces population densities in the food chain. Main branches could be seen as the main ‘trophic pathway’ [41], but upon any addition of a species, the main pathways can shift to another branch.

When following the build-up in Fig. 3a parallel branches with large biomass are possible (e.g. Fig. 3a, addition 6). Thereby the total biomass may substantially exceed that of a single initial species (Fig. 3c and Fig. 3d). The food webs obtained are however constrained by the overall decay compared with the total input from the basal level. Total species richness is also limited by the exact geometry of the food web. Food webs with many branching points often saturate earlier than those with predominantly parallel branches. Fig. 5 presents the average species richness of the largest food webs we obtained when no new additions were possible, i.e. when any addition would have violated the assembly rules or the sustainability conditions. We see that large *β* ≈ 1 and small minimal decay rates *α* allow for food webs with species richness on the order of 1/*α_min_*. Thus it is possible to reach realistic species richness (≈ 100 in the webs compiled by Dunne et al. [33]).

**Figure 5:**
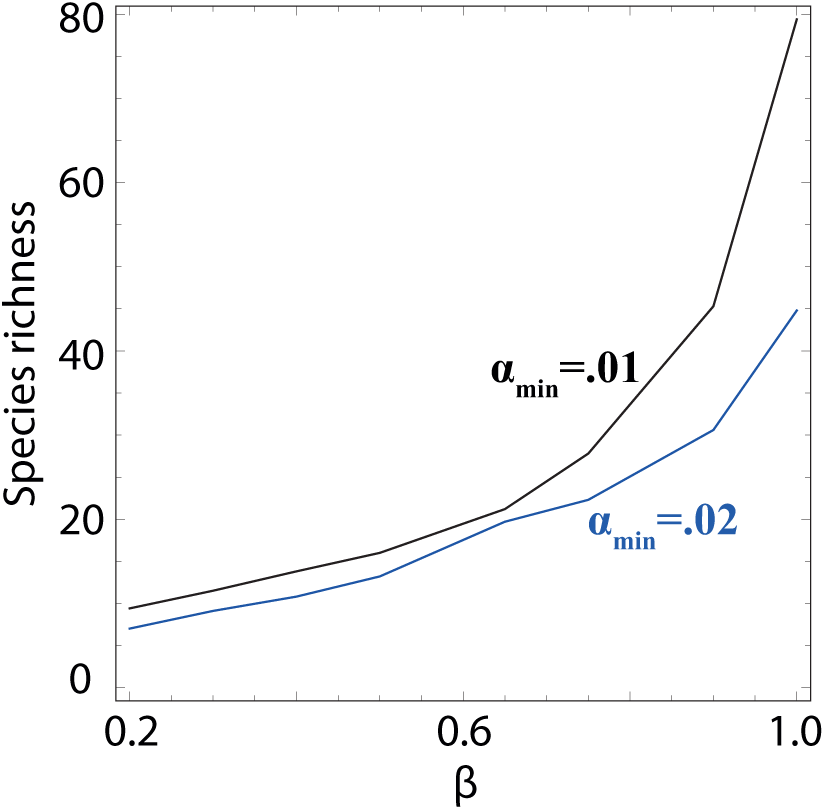
Total species richness obtained in assembly of tree like food webs. In our simulations, a single nutrient source was taken to provide the carrying capacity unity to trophic level one in the food web. Species richness depends on the minimal allowed spontaneous decay rate *α_min_*, two examples are shown for *α_min_* = .01 and *α_min_* = .02 (blue). The parameter *β* quantifies consumption efficiency, i.e. the proportion of consumed biomass that is available to the consumer in terms of population growth.

To put the decay rate parameter *α* into perspective it should be compared to the assumed maximal growth rate of 1. Therefore, for a species with a value *α* = 0.02 this corresponds to the ability of an individual to generate 50 descendants within its lifetime set by natural (non predatory) death of the individual. From Fig. 3 one also sees that the consumption efficiency *β* constrains maximally obtained species richness. However, even with low values of *β* a single nutrient source can support on the order of 10 species. Thus, when considering that real ecosystems can have many different nutrient sources, we conclude that food webs of realistic size can exist under well mixed conditions.

### Species richness in model and field data

A generic feature of our networks is that species richness is largest at intermediate trophic levels, while there are few basal or top predator species [26]. This is commensurate with data, e.g. from terrestrial [42] and marine food webs [43]. It is also in line with results from a semi-analytical effective model [44], where the interaction matrix was parameterized to yield inequalities for the means and variances of species concentrations. Also there it was found that intermediate-level species richness should dominate. A value of *β* = 1 thereby means perfect efficiency (Fig. 4a) whereas *β* = 0.2 (Fig. 4b) means imperfect efficiency, i.e. most of the energy is lost at every consumption step and is thereby not available to the consumer. In both panels *α_min_* = 0.01, but results are very similar when using *α_min_* = 0.02. Fig. 4c shows the empirically measured distribution of species richness versus trophic levels for seven different food webs. We see that our assembly process produces quite reasonable distributions, in spite of the fact that our model so blatantly disregards both the evolutionary process of removal of species, omnivory, cooperation as well as spatial effects and non-linear responses.

### The effect of additional weak links

Also, our model considers food webs with only a minimal number of strong links, implicitly assuming that most detected links are in fact weak. This assumption is certainly an idealization, however, we recall that empirical work suggests bi-modal link strength distributions, with few strong, but many weak links [16, 17, 18] — lending at least qualitative support to our assumption. To add realism, we augment our food webs by additional weak links between all species (example in Fig. 6). We find that as long as the strength of these additional links is kept below the value of *α_min_* feasibility is usually not threatened. In work on dynamical stability it was shown that for generalized Lotka-Volterra equations the equilibrium point and dynamical stability could be maintained by making simultaneous changes to the off-diagonal elements of the interaction matrix as well as the species growth rates and carrying capacities [14]. This shows that, once a feasible and stable steady state is obtained, nonzero off-diagonal terms could be “morphed”. Applying this to our weak links could allow these values to be further modified in a systematic way. Our results hence show that realistic food webs of arbitrary connectivity can, in principle, be sustainable.

**Figure 6:**
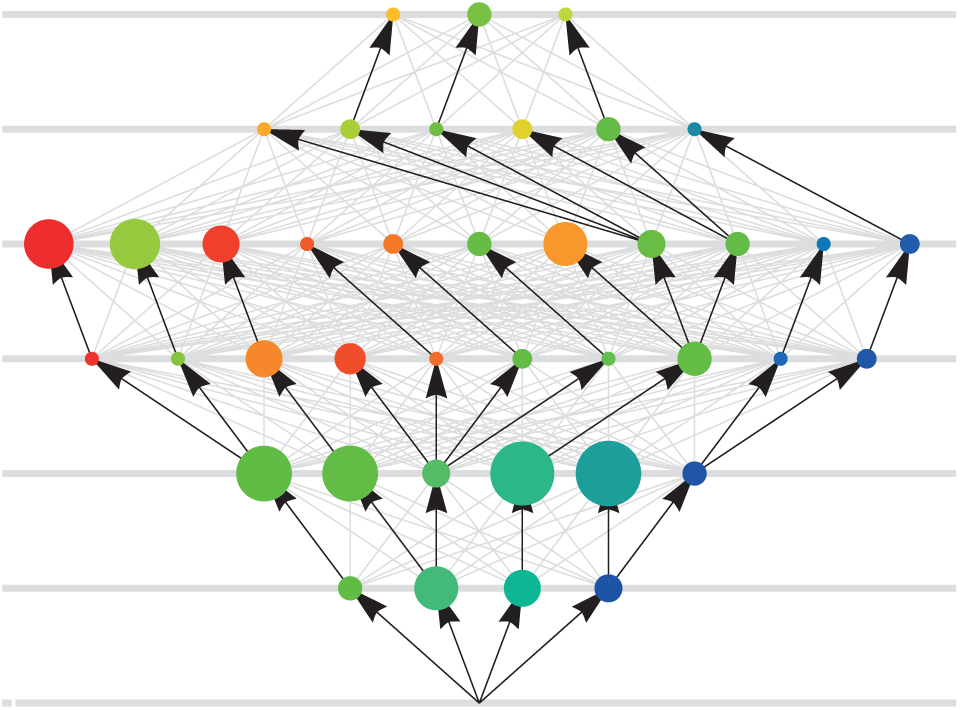
Food web showing species population densities. Species densities (circles) and feeding links (arrows). Circle areas are proportional to species densities. Colors from blue to red indicate earlier and later additions of species.

## Discussion

It is clear that large food webs are far from randomly connected. This is corroborated by both R. May's seminal work as well as subsequent empirical and theoretical studies. Basic rules for possible sustainable structures of food webs have been constructed by building on either the competitive exclusion principle [26, 29, 45, 46] or by constraining the consumer-resource couplings by saturating [25, 47, 48] or saturated couplings [49]. Within the simple generalized Lotka-Volterra equations the present work shows that for any food web with invertible interaction matrix parameters exist where all species can have feasible and dynamically stable population densities. We further find that this can even be obtained under the further condition that all interaction strengths be equal and only decay rates vary. The present paper specifies how these rates can be chosen so that stability and feasibility of the food web are maintained at each step during its assembly.

Using our procedure we easily obtained food webs with 5 trophic levels, and maximal species richness at levels 2 or 3. This distribution resembles the overall pattern found in real food webs with approximately 100 species (compiled by Dunne et al. [33]). Our analysis focused on one limiting nutrient source, whereas biodiversity in natural systems will likely be larger as the number of distinct sources is much greater than one. For our tree-like webs, separate or segregated resources would simply lead to an additive effect on species richness, with independent trees thriving separately, but would have no impact on the distributions of species richness we obtained. One implication is that large food webs in principle can have a total biomass that is substantially larger than the carrying capacity imposed on species at the basal level of the food chain.

Our study was constrained to the simplest, linear mass-action kinetics between consumer and resource, namely the type-I response. This simplification facilitated the clear division into *free* and *controlled* species. Our aim was to show that even within this constrained framework, where diversity is made more difficult, stable coexistence of many species at many trophic levels is possible. If one were to include type-II saturating response links, these inhibited interactions could prevent a consumer to completely control its resource which in turn can increase the parameter space for which many species could co-exist. It is thus remarkable that our species-richness distributions remain close to both the ones observed in the field and those assembled by using more complicated types of functional responses [50]. Notably, our present work further does not consider omnivory and parasitism, or self-interaction [51] and spatial dispersal [52, 53]. All these mechanisms can be alternative avenues to achieve coexistence, as can e.g. be seen by the current debate on the role of parasitism [26, 33, 36, 54, 55, 56].

Finally we emphasize that the current study focuses on monotonic assembly of food-webs, and thus entirely disregards the fact that invasion by new species will often force others to be eliminated. Such non-monotonic growth may bring about more fine-tuned selection of parameters for survivors, but will also render the long-term growth dynamics of the food web punctuated. At rare occasions, an invading fast growing basal species may even eliminate all species feeding on the same nutrients. Further studies of such evolutionary dynamics will be explored in future work.

## Acknowledgments

The authors acknowledge financial support by the Danish National Research Foundation through the Center for Models of Life.

This work was supported by a research grant (13168) from VILLUM FONDEN.

## Author Contributions

All authors have jointly developed the research ideas, worked out the equations and written the manuscript.

